# Coding of time with non-linear mixed selectivity in Prefrontal Cortex ensembles

**DOI:** 10.1101/2023.04.07.535754

**Authors:** Catherine A. Mikkelsen, Stephen J. Charczynski, Scott K. Brincat, Melissa R. Warden, Earl K. Miller, Marc W. Howard

## Abstract

Previous work has identified stimulus specific time cells as a potential mechanism for working memory maintenance. It has been proposed that populations of stimulus specific sequences of cells could support memory for many items in a list over long periods of time. This would require information about one stimulus to persist after the presentation of subsequent stimuli. However, it is not known if sequences triggered by one stimulus persist past the presentation of additional stimuli. It is possible that each new stimulus terminates preceding sequences, making memory for multiple stimuli impossible. To investigate this question, we utilized a data set originally published by (Warden & Miller, 2010), studying the firing of monkey prefrontal neurons during short lists of stimuli. We were able to decode “what happened when” throughout the list, using linear discriminant analysis. Additionally, we were able to decode the first stimulus after the presentation of the second stimulus. Furthermore, we found that stimulus modulated sequences of cells, with discrete temporal fields, continue after the second item was presented. Much of the information about the previous item was carried by neurons that responded to conjunctions of stimuli and the timing of late-firing cells was synchronized to the firing of the second stimulus rather than the first. These properties falsify a simple linear model of sequential time cells. These results suggest that non-linear mixed selectivity extends to continuous variables such as time, but that in this experiment at least, only the timing of the most recent stimulus was explicitly maintained in ongoing firing.

## Introduction

The prefrontal cortex has long been implicated in working memory (Fuster & Alexander, 1971; Goldman-Rakic, 1995; Jacobsen, 1936; Miller & Cohen, 2001). Cognitive models of working memory require multiple past stimuli to be represented in working memory (Atkinson & Shiffrin, 1968; Baddeley & Hitch, 1974); this is necessary to allow associations to be formed between stimuli for later recall (Kahana, 1996). Although early work focused on content-specific persistent firing of neurons in PFC as a neural model for working memory (Funahashi et al., 1989; Miller et al., 1991), more recent neurophysiological work studying firing of PFC neurons has identified more subtle and complex signatures of firing associated with maintenance of information in working memory. This paper focuses on how information about multiple past items, and the time at which they were presented, are expressed in the firing rates of neurons in PFC using a canonical dataset (Warden & Miller, 2010).

### Non-linear mixed selectivity in PFC ensembles

Consider stimuli that differ on multiple dimensions. For concreteness, let us consider two visual shapes, square or circle, and two different colors, red and blue. Content-specific firing for these shapes could be divided among the different features. For instance, some neurons might fire to circles, regardless of their color, while other neurons fire to squares, regardless of color. Another separate population could fire only to the color of the stimuli, and ignore the shape. In contrast, one could imagine a population where many neurons fire to conjunctions of stimuli. For instance, one neuron might fire only for red circles and fire for neither red squares nor blue circles. This form of coding is referred to as non-linear mixed selectivity. Rigotti et al., (2013) observed that firing in the PFC during maintenance of information in working memory for short lists included many conjunctive neurons that coded for features extended in time, such as neurons that coded for a particular stimulus in a particular serial position. Non-linear mixed selectivity endows a system with many computational advantages, at the cost of requiring many more neurons to cover the stimulus space than would have been necessary for neurons that only coded for one or another feature (Fusi et al., 2016; Sreenivasan & D’Esposito, 2019). Mixed selectivity in the prefrontal cortex may also be a reflection of the top-down influence of attention on primary cortices (Sreenivasan et al., 2014). The utility of non-linear mixed selectivity is supported by the finding that conjunctive coding is also observed in the hippocampus, for a wide range of variables (Anderson & Jeffery, 2003; Komorowski et al., 2009; Nieh et al., 2021; E R Wood et al., 2000; Emma R Wood et al., 1999). Indeed, the fact that hippocampal place cells exhibit radial basis functions for position has long been argued to reflect conjunctive coding for distance to different landmarks available in an environment (O’Keefe & Burgess, 1996).

### Timing information in PFC ensembles

Representations of the time at which items were experienced has long been an important feature of cognitive models of working memory (Brown et al., 2000; Hacker, 1980; Howard et al., 2015). In the last decade, a growing body of work has demonstrated that the firing of neurons in a wide range of brain regions carry information about the time at which past events were experienced (MacDonald et al., 2011; Mello et al., 2015; Rossi-Pool et al., 2019; Tsao et al., 2018). In the hippocampus, sequentially-activated time cells (MacDonald et al., 2011; Pastalkova et al., 2008) can be used to reconstruct the time since a delay interval began. Because different stimuli in a working memory experiment trigger distinct sequences, hippocampal time cells can be understood as conjunctively coding for what happened when in the past (Cruzado et al., 2020; Taxidis et al., 2020; Terada et al., 2017). The same type of conjunctive coding of what happened when in the past can be observed in sequentially-firing neurons in PFC (Cruzado et al., 2020; Tiganj et al., 2018). Sequentially-activated time cells are not the only way a neural population may code for the time of past events. For instance, in entorhinal cortex, time can be decoded from populations of neurons that change their firing monotonically over time, but at a wide variety of rates (Bright et al., 2020; Tsao et al., 2018). Indeed, many authors have noted that the firing of PFC neurons carries information about time while not explicitly describing sequential firing (Cavanagh et al., 2018; Cueva et al., 2020; Murray et al., 2017).

### Temporal mixed selectivity in lists of multiple items

Prior work has established that prefrontal neurons use non-linear mixed selectivity to code for information maintained in working memory after a brief list. Prior work has also established that sequentially-activated neurons code for time by tiling the delay following a stimulus, showing conjunctive coding of what happened when. However, previous work on timing information has focused on delays following a single item (but see Goh, 2022). This leads to a critical question. As the number of items in a list grows, the number of neurons necessary to conjunctively code for the list grows exponentially (the number of lists composed of *N* items of length *L* goes like *N*^*L*^). This concern becomes more acute if the ensemble also retains information about the time of each of the separate items in the list. In this paper we study the retention of information in working memory during study of short lists of visual stimuli using an existing dataset (Warden & Miller, 2010). The primary question is how information about the time and identity of earlier items is retained following the presentation of the second item in the list.

## Methods

### Description of Behavioral Task

Warden & Miller (2010) trained two monkeys to remember lists of two visual stimuli. Each of the four clearly distinguishable images could appear in either list position. The same stimulus could not appear twice in the same list. Each stimulus was presented for 500ms with a 1s delay after each presentation. For this paper, only the time of list presentation, or the first 3000ms of each trial, was considered. After each list, the animals were presented with one of two behavioral decision tasks that required the monkey to remember the list. The data from both tasks was combined. Recordings were made in the prefrontal cortex. See Warden and Miller (2010) for complete methods.

### Population Analyses

Our primary interest is to know how the firing of PFC neurons after the second list item was presented reflected memory for the first list item. To evaluate this, we will attempt to decode the stimulus in the first serial position at all points during presentation of the list. As a control, we will attempt to decode the stimulus in the second serial position at all time points during list presentation. Decoding accuracy of the second list item prior to the time it was presented provides a baseline that controls for dependencies between the serial positions and any other methodological issues that may arise from the decoder.

### Linear Discriminant Analysis Serves as the Base of the Decoding Analyses

A cross temporal classifier was used to identify stimuli based on neural firing data. This involves running a linear discriminant analysis for all of the combinations of time bins. The classifier was trained to decode the stimulus on each trial using firing from a given time bin and then tested on firing from all time bins. The linear discriminant analysis uses firing rate averaged over each bin on each training trial to create a linear model, for which each neuron is a variable. Firing on a separate set of test trials is then evaluated by the model to predict the probability of the stimulus identities for the test trials. We implemented the classifier using the Matlab function “classify”. Random normally distributed noise was added to the training and test data with a sigma of 10^−7^ and a mu of 0, in order to prevent singularities.

### LDA is utilized to determine “what”

We sought to decode the identities of the stimuli presented in both presentation positions. Firing was divided into 250ms blocks. Each run contained 260 training trials, 40 test trials, and 200 units. Units were selected that have at least 420 trials (1.4 x (training trials + test trials)). 50 repetitions were run of this analysis and the average accuracies were calculated. We compared the bins representing equal offset for the classification of the second stimulus after the presentation of the first, and the classification of the first stimulus after the presentation of the second. For example, we compared classification of stimulus 1 at 2000 ms (500 ms after the other item was presented) to classification of stimulus 2 at 500ms (500 ms after the other item was presented).

### LDA is utilized to determine “when”

We also sought to decode time to determine if temporal information persists throughout the entirety of the trial. We again used a linear discriminant analysis, but divided the trial into ten 300ms bins. The classification of time bin used 200 training trials and 60 test trials. Units were included that had at least 312 trials (1.2 x (training trials + test trials)). We used 150 units instead of 200 due to the differences in number of trials needed for the increase in the number of outcomes.

### Analysis of Individual Cells

We sought to characterize the firing field of each neuron using a series of models that were fit to the spiking profile. We included four models: a constant model, a “pure time’’ model with Gaussian receptive fields, a stimulus-specific time cell model, and a conjunctive time cell model. Parameters for each model were selected to maximize the likelihood of the observed spike train across sequences. While the Gaussian time field model does not capture the nuances of the firing fields, the representation is sufficient to identify cells whose firing is modulated by the stimulus identity and temporal patterns.

First, we identify all time cells. To do this we compare the log likelihoods of the model fits for the Gaussian model to the constant model. Next the time cells are divided into three mutually exclusive subpopulations: pure time cells, stimulus-specific time cells, and conjunctive time cells. “Pure time’’ cells are those whose fit was not improved by adding parameters sensitive to the stimuli presented on each trial. The stimulus-specific model has four separate stimulus parameters in addition to the parameters of its time field. The conjunctive model has 12 parameters one for each possible list (recall that the four stimuli were never repeated within a list). The stimulus-specific time cells are defined as those for which the stimulus-specific model is a better fit than the Gaussian and conjunctive models. The conjunctive time cells are defined as those for which the conjunctive model is a better fit than the Gaussian and stimulus-specific models.

Model fits and parameter selection were performed on the cells using a customized program Maxlikespy (https://github.com/tcnlab/maxlikespy), which utilizes scipy’s basin-hopping method to determine model parameters. This method is similar to previous work (Cruzado et al., 2020; Tiganj et al., 2018). The basin hopping algorithm was run until a better fit could not be found for 1500 iterations. It uses the Truncated Newton (TCN) method as the minimization method. The first model simply estimated a constant firing rate:

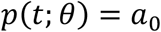

The ‘‘pure time’’ model adds a Gaussian temporal receptive field:

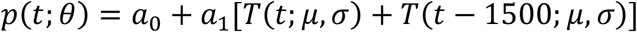

Where

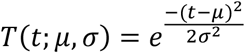

The stimulus specific time cell model allows us to consider the influence of stimulus specificity on the firing of the neurons in addition to temporal specificity. It considers each of the four stimuli separately. For the stimulus specific Gaussian model, the value of *C*_*i*1_ was set to one for trials when the stimulus was presented in the first location, and *C*_*i*2_ was set to one for trials when the stimulus was presented in the second location.

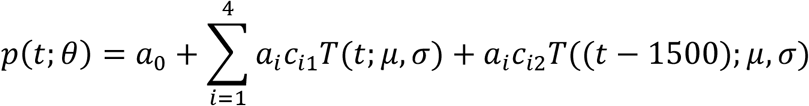

The conjunctive time cell model has separate information about the identities of the stimuli in the first and second presentations. For the conjunctive model the value of *C*_*i*_was set to one for trials when the associated list was presented (ie. both stimuli in order), and zero for all other lists.

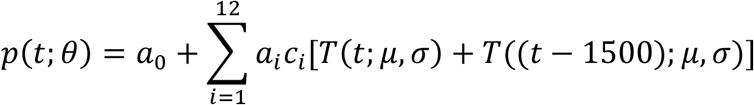

For all temporal models, the model fits are able to account for two peaks in a cell’s firing that are 1500 ms apart. The mu value was allowed to vary from 0 to 3000ms. Sigma is allowed to vary from 0.001 to 1000ms. The coefficients are bounded such that the sum of all a’s is less than 1. To correct for differing numbers of parameters we used the Matlab function “lratiotest” using a p-value of 0.01 (Bonferonni corrected). The constant model has one parameter, the pure time cell model has 4 parameters; the stimulus specific time cell model has 7 and the context-dependent stimulus specific time cell model has 15.

### Evaluating distributions of time cell parameters

Prior work studying time fields following presentation of one item shows a monotonic, roughly linear, relationship between time field width and the peak time. Inspection of analogous plots for this dataset showed an apparent discontinuity after the second list item was presented at 1500 ms. To evaluate this hypothesis, we compared a regression of parameters over values of mu spanning the entire duration of the list (0 to 3000 ms) to a piecewise regression that fit the intervals 0 to 1500 ms and 1500 ms to 3000 ms separately. Each of these regressions used constant and linear terms. We compared the best-fitting model over the entire interval to the best-fitting piecewise models using Akaike Information Criterion (AIC) and Bayes Information Criterion (BIC).

### Classification by Subpopulation

The stimulus classifier approach utilized for the entire population can be modified to be combined with the analyses of subpopulations of cells. For this analysis we analyzed only the bins for which training and testing are equivalent. Because of the decreased number of cells available, only 50 units were used per repetition. There were still 260 training trials and 40 test trials.

## Results

### Neural data can be used to classify “what” happened throughout the trial

*A* cross-temporal classifier was able to decode the identity of the first stimulus above chance even after the presentation of the second stimulus, as shown in Figure 1. The accuracy of the decoder decreases from 250ms until the presentation of the second stimulus (p < 0.01, slope= -2.5×10^−7^). While the ability of the classifier to decode the identity of the first stimulus drops off over time, it did not return to chance after the presentation of the second stimulus. The box plot in Figure 1d shows the accuracy of the decoding of the first stimulus (blue) for all time points across the diagonal, as compared to the accuracy of the decoding of the second stimulus (black), for all time points across the diagonal. We used two tailed two-sample t-tests on the distributions of the classifier accuracies for a given set of parameters to compare these two epochs. For example, we can compare the decoding of stimulus 2 from 0-250 ms to the decoding of stimulus 1 from 1500-1750ms. Repeating this process, we find that each of these pairs is significantly different (p<0.01). There are 98 degrees of freedom for all pairs. The t-statistics are, in order from stimulus onset, 11.9815, 9.7968, 10.2802, 11.5106, 10.3817, 12.5892.

**Figure 1:**
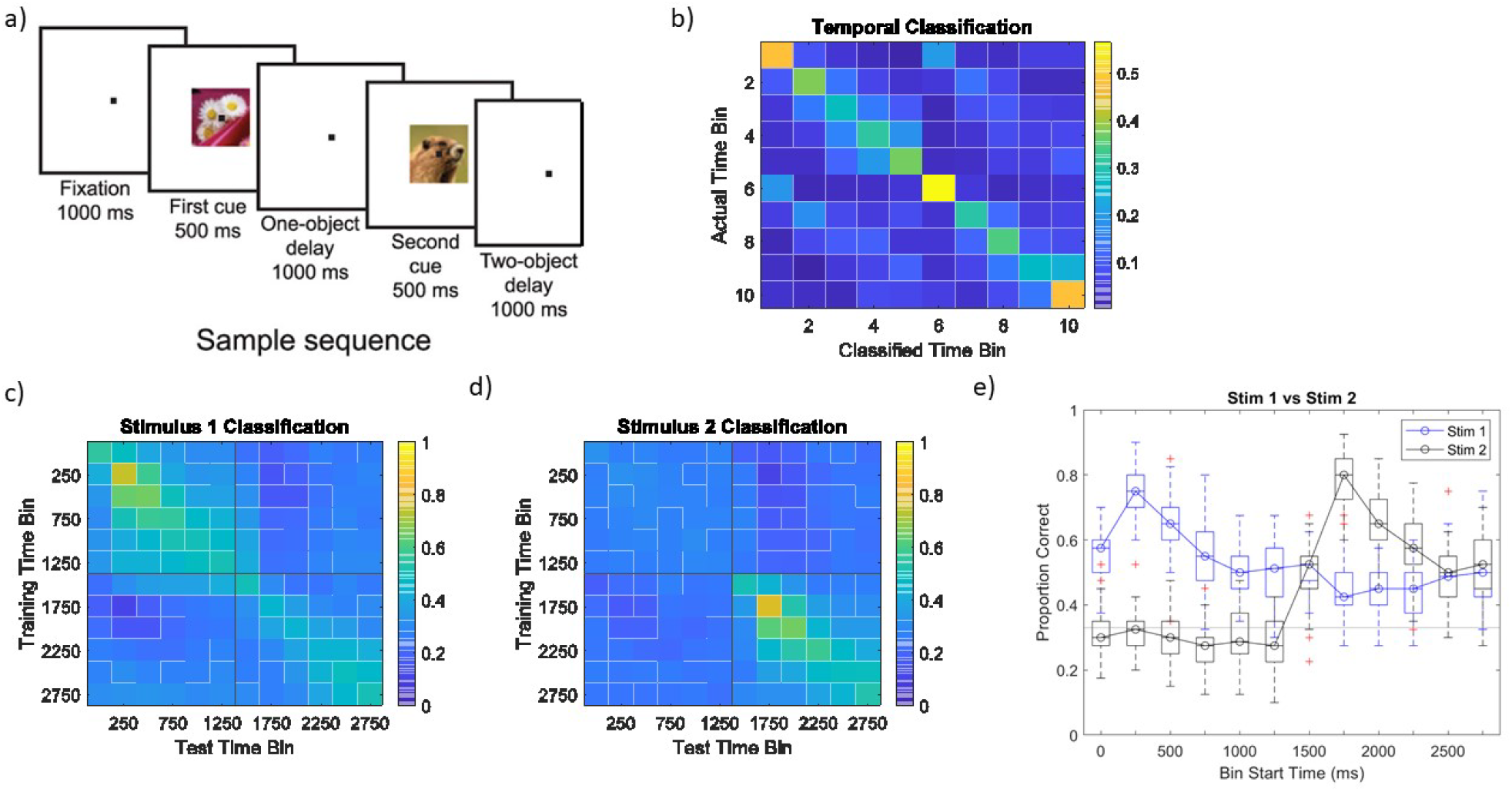
The population of neurons represents the identity and time of the first stimulus throughout the trial. a) A description of the task. The animal is presented with a stimulus from 0-500ms, followed by a delay from 500-1500ms. The second stimulus is presented from 1500-2000ms, followed by another delay from 2000-3000ms. b) A linear discriminant analysis (LDA) shows the ability to decode time bins. Each row represents the actual time bin for a set of decoded trials. The number of classified time bins for each row is counted and then normalize by the number of trials for the row. c) A cross temporal classifier is able to decode the identity of stimulus 1, even after the presentation of the second stimulus. d) A cross temporal classifier is able to decode the identity of the second stimulus, after it is presented at 1500ms. e) This graph shows the accuracy of the cross temporal classifier for when the training time bin and the test time bin are the same. The accuracy increases rapidly for stimulus 1 classification (blue) and then slowly drops off, but never to chance. The accuracy for stimulus 2 classification (black) increases after the presentation of the second stimulus at 1500ms.

### The neural data can be used to classify “when” throughout the trial

Time can also be decoded from the neural data (Fig 1b). This shows that there is temporal information carried in the neural firing. If we count the number of classifications for each potential time bin, for each row, the distribution is significantly different than for a uniform distribution (p<0.01). We found that the distribution of the probabilities was significantly different from chance for each row using 10-sample test for given proportions *x*^2^(10) ranging from 169.4 to 585.8, all p < .001.

### Temporal parameters from model-based analysis

The overwhelming majority of units, 792/867 were better fit by a model with a Gaussian time field than by the model with only constant firing rate. These 792 “time cells’’ were further classified as 189 pure time cells, 367 stimulus-specific time cells, and 214 conjunctive time cells based on criteria described in the methods. We describe the properties of each of these groups below.

### Pure Time Cells

Figure 2 summarizes the properties of the units classified as “pure time cells’’. These cells fire in a temporally modulated way irrespective of the stimulus presented. The heatmap in figure 2a shows that many of these cells have two fields, equally distant from each stimulus presentation. However, there exists a subpopulation of cells (see red circle) that fire only after the presentation of the second stimulus. These cells may also be sensitive to serial position, and could also be successfully fit by a model that combines time with presentation period. Alternatively, they could be triggered by the first item, but at a delay longer than 1500 ms. However, these two possibilities are confounded due to the equal spacing on each trial.

**Figure 2:**
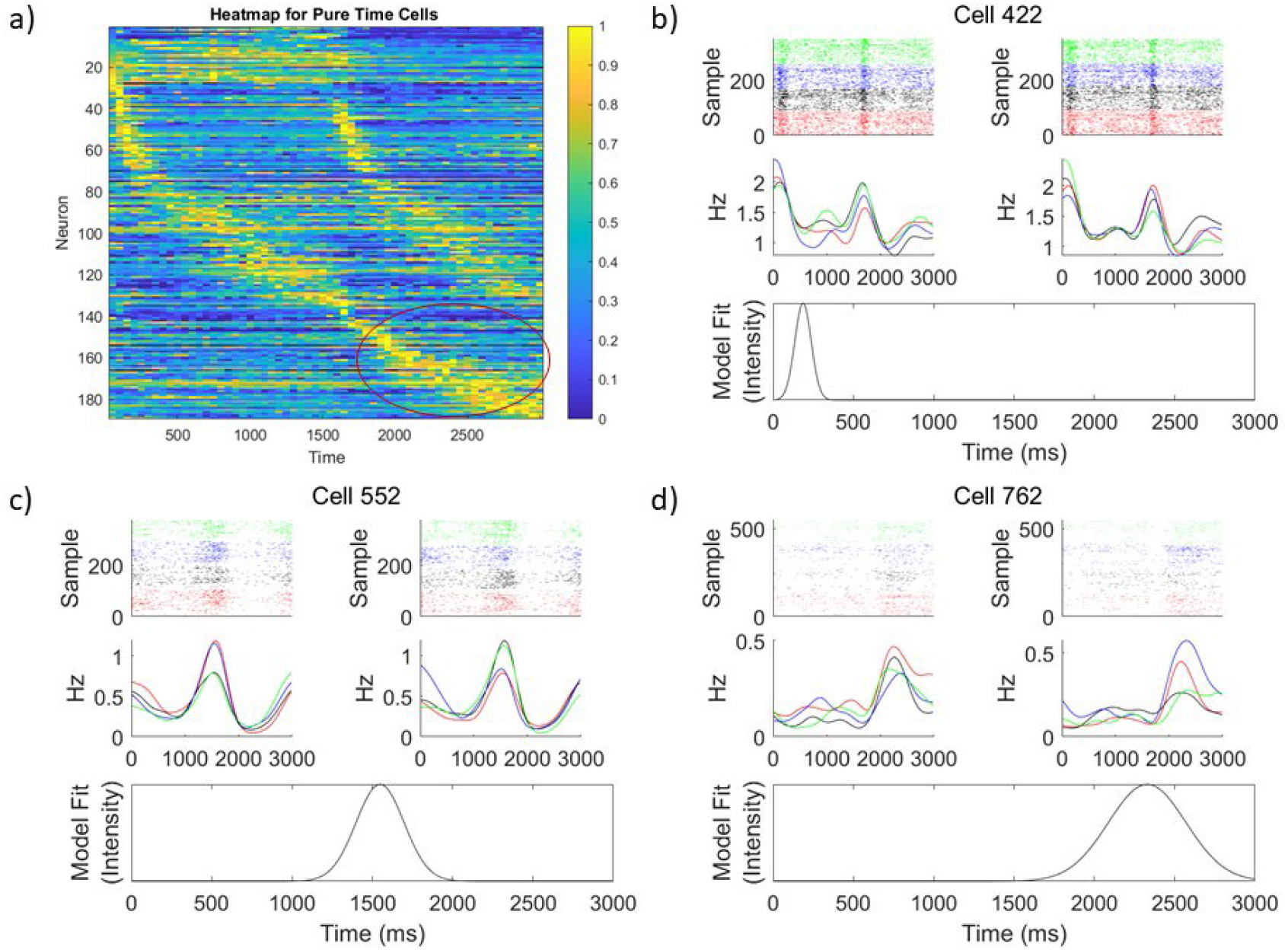
Single units best fit by the pure time model. a) A heat map of the pure time cells, sorted by the mu value of the Gaussian only model, shows cells that peak only after the presentation of the second stimulus. Each row represents the self-normalized firing of each neuron. Each cell is normalized to its maximum firing rate bin. b-d) Rasters of time cells show punctate fields. Samples are color-coded and sorted by stimulus 1 identity on the left, and stimulus 2 identity on the right (trials where stimulus A is shown are in red, stimulus B in black, stimulus C in blue, and stimulus D in green.) The top panel shows rasters where each row represents a trial, and each line marks times the cell fired within that trial. The second panel is the Gaussian smoothed firing rate, calculated separately for each stimulus identity. The bottom panel shows the model fit for the Gaussian model.

We see a diversity of sigma and mu values (Fig 5) for this subpopulation. Investigating the relationship between the mu and sigma values, the best model fit was the split linear model (ΔAIC=27.8, ΔBIC = 21.3). The fact that the piecewise linear model was a better fit argues against the hypothesis that the cells with mu > 1500 ms reflect linearly increasing sequences initiated by the first stimulus.

### Stimulus-Specific Time Cells

Stimulus specific time cells are the 367 cells for which the cell’s firing is modulated by time and the stimulus-identity. Stimulus-specific time cells also tile the entirety of the trial, and cells responsive to a stimulus presented first may have temporal fields after the presentation of the second stimulus. Shown in figure 3a, we note that there is a subpopulation of 31 cells that fire to a particular stimulus at a time greater than 1500ms (8.4% of stimulus selective cells), or after subsequent stimuli have been presented (red circle). Early and mid-trial examples of stimulus-specific time cells are shown in Figure 3b and 3c. Figure 3d shows an example of a stimulus-specific time cell with a late temporal field. Importantly, the heatmap appears to terminate its “hook” at 1500ms.

**Figure 3:**
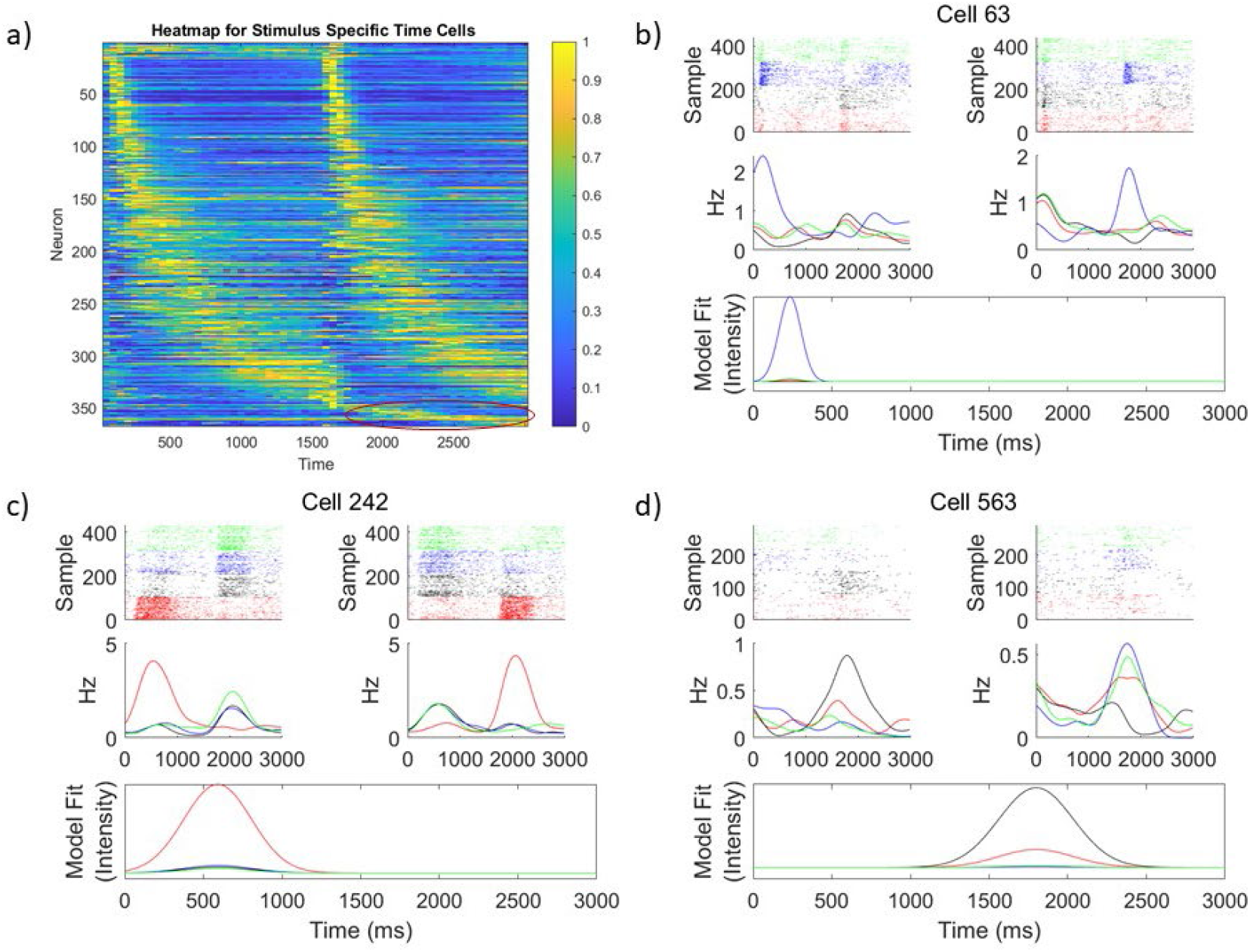
Single units best fit by the stimulus selective time cell model. a) A heat map shows that the stimulus-specific time cells tile the entirety of the delay with punctate fields. Each row represents the normalized firing of a neuron, and neurons are sorted by the model mu value of the conjunctive model fit. b-d) Examples of rasters for stimulus specific cells show punctate fields as well as variations in firing dependent upon the stimulus presented for that trial. Samples are color-coded by and sorted by stimulus 1 identity on the left, and stimulus 2 identity on the right (trials where stimulus A are shown are in red, stimulus B in black, stimulus C in blue, and stimulus D in green.) The top panel shows rasters where each row represents a trial, and each line marks times the cell fired within that trial. The second panel is the Gaussian smoothed firing rate, calculated separately for each stimulus identity. The bottom panel shows the model fit of the stimulus specific model. This model shows firing for stimulus A in red, stimulus B in black, stimulus C in blue, and stimulus D in green.

Additionally, there is a discontinuity in the sigma and mu values between the first and second presentation periods (Fig 5), as shown by comparing a piecewise linear model associating the sigma and mu values to a linear model, (ΔAIC= 45.8 and ΔBIC=38.0). This fails to support the hypothesis that in this experiment mu and sigma increase linearly with time. Further experiments, dissociating ordinal and continuous time, and model testing would be required to confidently determine if the cells are firing to stimulus 1 at a late time, or stimulus 2 but only in the second presentation period.

### Conjunctive Time Cells

Conjunctive time cells are cells that fire to a specific list presentation, or combination of a specific pairs of stimulus 1 and 2 identities (ex. C followed by A). Figure 4a shows a heat map of these cells. Many of the conjunctive cells fire after the presentation of the second stimulus. These cells also encode information about the first stimulus after the presentation of the second stimulus, because they fire to a specific pair of stimuli. Ordinal position is also explicitly encoded in the list pair. Figures 3 b-d show examples of these cells at early (b), middle (c), and late (d) times within the trial. We also investigated the relationship between the mu and sigma values for the conjunctive cell population. For conjunctive time cells the linear and piecewise models were very similar, and the best fit model differed based on the criterion used (ΔAIC= 0.5, and ΔBIC= -6.2).

**Figure 4:**
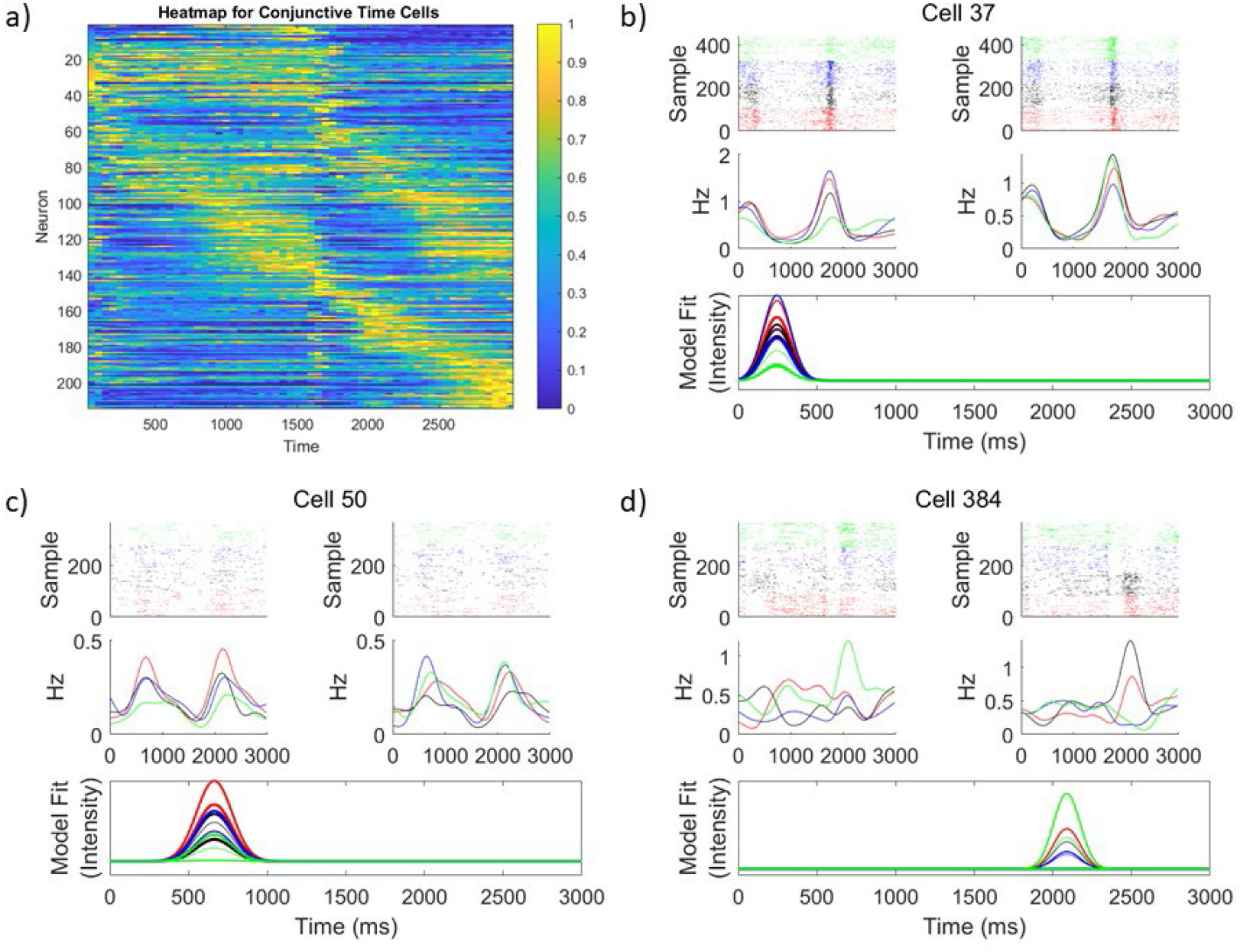
Single units best fit by the conjunctive time cell model. a) A heat map shows that the conjunctive time cells, which are selective to a particular combination of 1st and 2nd stimulus identities, tile the entirety of the delay with punctate fields. Each row represents the normalized firing of a neuron. Neurons are sorted by the model mu value of the conjunctive model fit. b-d) Examples of rasters for conjunctive cells show variation in firing over time. Samples are color-coded by and sorted by stimulus 1 identity on the left, and stimulus 2 identity on the right (trials where stimulus A are shown are in red, stimulus B in black, stimulus C in blue, and stimulus D in green.) The top panel shows rasters where each row represents a trial, and each line marks times the cell fired within that trial. The second panel is the Gaussian smoothed firing rate, calculated separately for each stimulus identity. The bottom panel shows the model fit of the conjunctive model. In this graph the colors of the lines represent the identity of the first stimulus (red= A, black=B, blue=C, green = D), and the width of the line represents the identity of the second stimulus (moving from thinnest to thickest from A to D). For example, a model that predicts activity for the condition of stimulus B presented first followed by stimulus A would be represented by a thin black line.

### All subpopulations can classify stimulus information independently

In order to determine the source of the stimulus one information that persists past the presentation of the second stimulus, we combined the individual cell analyses with the decoders. Utilizing the three distinct subpopulations of cells: pure time cells, stimulus-specific time cells, and conjunctive time cells, we used the stimulus classifier on each cell type independently. Comparing results for cells with equivalent training and test times, shows significant differences for the ability to classify the identity of the first stimulus identity during the second presentation period, as compared to the ability to classify the identity of the second stimulus during the first presentation time, for time bins 4 and 6 (p < 0.01), for the pure time cells. For the stimulus specific and conjunctive subpopulations, the classification of the first stimulus during the second presentation period was better than the classification of the second stimulus during the first presentation period was significantly better (p<0.01) for all 6 time bins. The full characteristics of the classifier can be seen in Figure 6.

## Discussion

During presentation of the second item in the list, the firing of ensembles in PFC carried information about what happened when at both positions during the list. The identity of the first item from the list could be robustly decoded throughout the entire list (Fig 1). The time within the study list could also be decoded (Fig 1). However, because the time between the first and second item was fixed across trials it is not possible to distinguish timing information of the first item after the second item was presented. Analyses of single units showed that the population carried information about time via consistent firing profiles. The pure time model with Gaussian temporal firing fields better fit the data than a constant model for the overwhelming majority of cells (792/867). Although some neurons showed temporal profiles that were more complex than Gaussian receptive fields (see Fig 2,3,4), nonetheless the population demonstrated sequential firing, in that different neurons fire reliably at distinct points in the delay interval. This can also be seen by noting that the mu parameter, describing the peak time of firing, took a wide range of values across units.

Notably, stimulus information and temporal information were intimately related. This can be seen from the cross-temporal classifier, which showed a strong dynamic profile (Fig 2) and the broad distribution of temporal parameters for each of the subpopulations of neurons (Fig 5).

**Figure 5:**
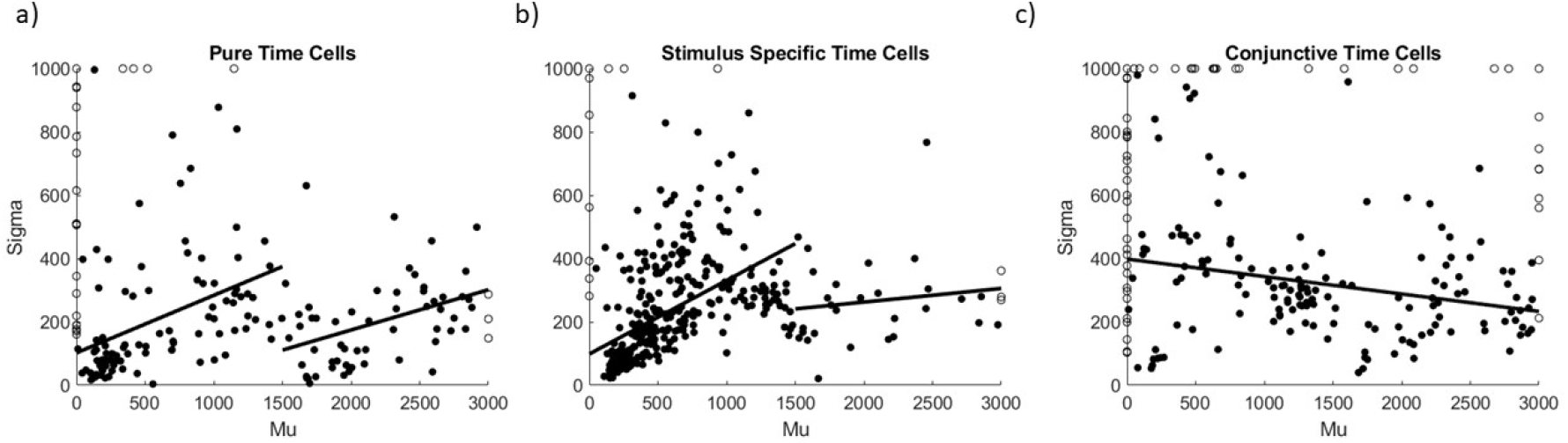
The relationship between the time field peak and width is non-linear. Scatter plots of the mu and sigma values of the model fits show the relationship between the time field peak time and the width, for all three subpopulations: a) pure time cells, b) stimulus specific time cells, and c) conjunctive time cells. In a and b, black lines show split linear models for two temporal epochs: the first spanning the first stimulus presentation period from 0-1500ms, the second spanning the second stimulus presentation period from 1500-3000ms. In c, the linear model is displayed showing the linear model extending from 0-3000ms.The unfilled black circles show cells with model fit values at a boundary; these weren’t included in the regression analyses.

### Stimulus information

The population showed robust stimulus coding for the most recently-presented item throughout the duration of list presentation (Fig 1). To assess the form of stimulus coding we constructed three subpopulations of cells. These subpopulations are a convenience for understanding the data that depend on choices of the experimenter; they should not be taken to be an argument that there are distinct biologically-meaningful categories of cells. Nonetheless, these artificial subpopulations revealed several properties of the ensemble coding.

Consistent with previous analyses of these data (Rigotti et al., 2013), the single unit analysis showed many neurons that code for conjunctions of list items (the conjunctive subpopulation Fig. 4). In addition, there was also a substantial number of neurons that showed simple coding for the most recently-presented item (the stimulus-specific subpopulation, Fig. 3). The “pure time’’ subpopulation, by construction, showed less stimulus coding than the other two subpopulations (by choosing a more stringent threshold between the populations we could have turned down the amount of residual stimulus information in the pure time subpopulation).

The subset of the pure time subpopulation that peaked after 1500 ms suggests that some neurons may be sensitive to serial position in addition to conjunctions of items. Notably, because the identity of the first list item was able to be decoded after 1500 ms from the conjunctive subpopulation, these neurons cannot simply be responding to serial position X time X current item.

### Temporal information

As in many previous studies (Bright et al., 2020; Cruzado et al., 2020; Kraus et al., 2013; Zoran Tiganj et al., 2018), the accuracy of temporal information decayed in precision with the passage of time since the most recently presented item. This can be seen by noting the decrease in the accuracy of the time decoder (Fig 1) and in the curvature of the heatmaps (Figs. 2,3, and 4). In the time following presentation of an item these plots show a reflected-J pattern. This indicates that the unfolding of the sequence “slows’’ with the passage of physical time in that the number of cells that begin firing at a time T after the stimulus goes down with T. This result is broadly consistent with the decrease in the spanning dimension of a neural ensemble with the passage of time (Cueva et al., 2020). The increase in the spread of firing fields with mu (Fig 5) is also consistent with a decrease in temporal accuracy with the passage of time. Both of these properties are characteristic of time cells described in PFC (Cruzado et al., 2020; Zoran Tiganj et al., 2018) and other brain regions (Cao, Bladon et al., 2021).

However, unlike previous findings, there appears to be a discontinuity in the sequences triggered by the first list item when the second list item was presented. This can be seen as a vertical line around 1500 ms in the heatmaps (Figs 2, 3, and 4) and as an apparent discontinuity in the sigma/mu plots (Fig 5). One way to understand this finding is that the presentation of the second item in the list not only initiates a new sequence, but disrupts the ongoing sequence. The identity of the previous item is preserved in the ensemble, largely due to conjunctive coding (Fig 6) but the sequence it triggered is terminated. It is unclear whether these results would also hold if the timing between the first and second item was variable across trials or if it was important to perform the task. There is good evidence that the timing of preceding events is preserved in other brain regions and other tasks. For instance, Tsao et al. (2018) showed that ramping neurons in rodent LEC triggered by the initiation of a session in an open field persisted throughout the session, despite periodic contextual changes. Recording from rodent hippocampus, Shahbaba et al. (2022) showed long sequences of firing during study of a five item list and were able to decode time within the entire list. However, because the list was consistent across trials, it is unclear whether sequences from early items were terminated or not by subsequent items. Future work should explore how multiple past events that vary in their identity and timing are represented in working memory in various brain regions.

**Figure 6:**
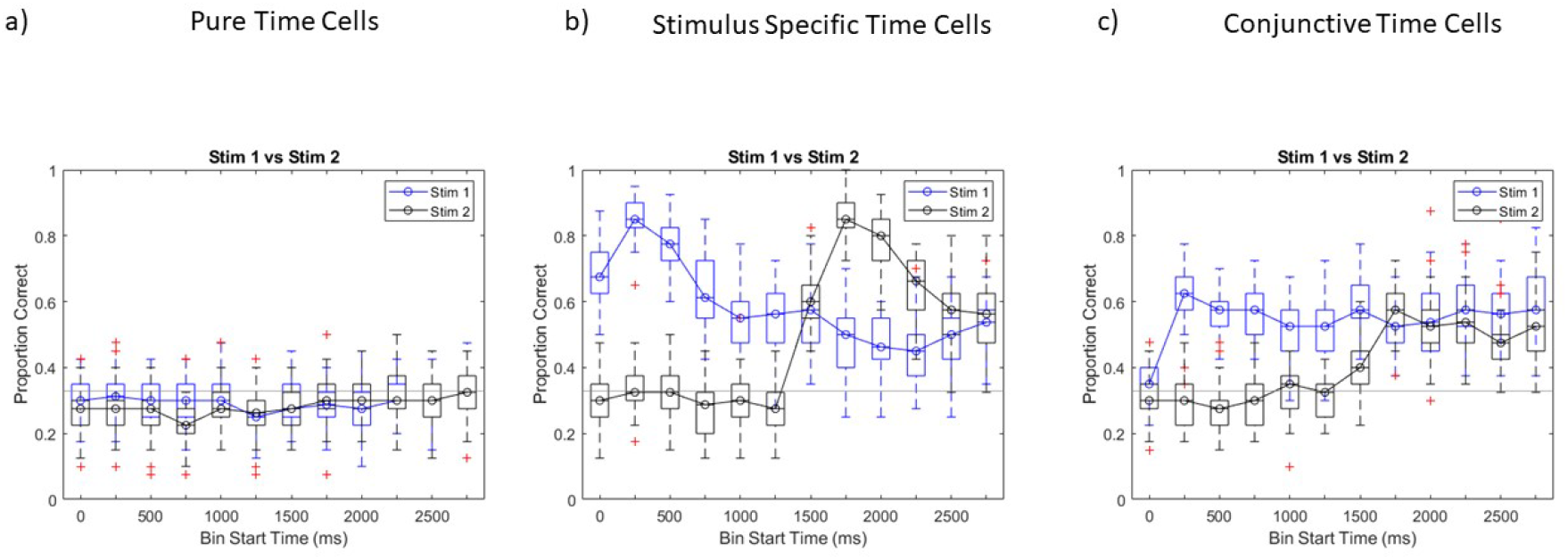
All cell categories can distinguish the identity of the first stimulus after the presentation of the second. Utilizing the stimulus classifier on a) pure time cells, b) stimulus specific time cells and c) conjunctive time cells, the ability to decode the stimulus 1 and stimulus 2 identity was examined at all time points. Boxplots represent the median, and interquartile range of the decoding accuracies. Blue represents decoding of the first stimulus. Black represents decoding of the second stimulus. Significant (p < 0.01) stimulus 1 information is available after the presentation of stimulus 2 at 1500ms at all 6 delays for the stimulus specific and conjunctive subpopulations. The classifier is only significant for the 4^th^ and 6^th^ time bins for the pure time cell population.

## Acknowledgments

Supported by ONR MURI N00014-16-1-2832 (Michael E. Hasselmo, PI). Office of Naval Research N00014-22-1-2453, The JPB Foundation, and The Picower Institute for Learning and Memory

